# Background proteome correction promotes confident identification of dynamic protein-protein interactions between different biological contexts

**DOI:** 10.64898/2026.09.01.748607

**Authors:** Melina A. Brunelli, Lisa Morishita-Cartwright, Natalie M. Clark, D.R. Mani, Samuel A. Myers

## Abstract

Affinity purification-mass spectrometry (AP-MS) enables the characterization of protein-protein interactions (PPIs), and the ease and sensitivity of such experiments has progressively increased. Beyond steady-state interactions of target proteins, a strong interest has emerged in monitoring how PPIs change upon significant biological perturbations, such as in disease contexts or small molecule modulation of the target protein. These perturbations likely not only induce PPI changes but can also lead to altered expression of proteins not of direct interest. Changes in protein abundance may alter which proteins adsorb to the affinity purification matrix, and due to the sensitivity of modern mass spectrometers, these differential “background binders” can masquerade as differential interactors. Contemporary approaches often do not account for differences in the background proteome, potentially inflating the number of false positives and negatives reported. Here, we provide technical considerations for the reliable annotation of dynamic PPIs, using the O-GlcNAc transferase (OGT) as a case study. We describe the installation of affinity epitope tags on endogenous OGT in mouse embryonic stem cells (mESCs), which we then apply for OGT interactor identification via AP-MS. We show that accurate representation of the bead background, which depends on the affinity matrix in use, is critical for elimination of false positive and false negative PPIs. This became even more pertinent as OGT PPI dynamics were measured under OGT catalytic inhibition via OSMI-4, which is known to perturb gene expression. The proteomes of OSMI-4-treated and control-treated mESCs differed, leading to distinct bead backgrounds in which the differential background proteins appeared as interaction gains or losses. These false positives were resolved by incorporating straightforward experimental controls through a practical statistical framework, allowing for a direct and confident comparison between treatment conditions. Incorporating these considerations into workflows investigating PPI dynamics will improve data fidelity and reproducibility.

## Introduction

The cellular proteome requires intricate organization for tight and dynamic coordination of cell signaling programs. One way this occurs is through physical interactions between proteins, namely protein-protein interactions (PPIs). Affinity purification-mass spectrometry (AP-MS) is the method of choice for the characterization of novel PPIs (1–4), with its accessibility and reliability scaling alongside advances in protein enrichment tools (5,6), mass spectrometry hardware (7,8) and data analysis tools (9,10). With this, it has become increasingly desired to measure PPI dynamics across different biological conditions by AP-MS. This could include comparisons of drug or small molecule modulation of target proteins (11), introduction of relevant clinical mutations (12), or comparison of healthy versus diseased states (13–15). The identification of differential PPIs across biological states may reveal novel targets for therapeutic development (16).

A confounding variable in these studies is that the biological contexts of interest likely harbor dramatically different cellular proteomes, which can affect one’s ability to discern changes in *bona fide* interaction partners from changes in non-interacting proteins (*i.e.*, “background binders”) adsorbing to the enrichment matrix (*i.e.*, “bead background”). As a result of the sensitivity of modern mass spectrometers, background binders are readily identifiable and can be misinterpreted as specifically enriched interactors. For example, overexpression of an epitope-tagged transcription factor for AP-MS characterization may result in gene expression–and ultimately, protein abundance–changes, relative to an empty vector or GFP expression vector control. Proteins now exclusively induced by transcription factor overexpression may be quantitatively, if not qualitatively, enriched in the bead background. This will cause these background binders to appear as enriched with the target protein, regardless of their true interaction status. This challenge compounds as more proteome-altering variables are included in the experimental design (*e.g.*, cancer vs. healthy tissue, drug treatment vs. vehicle). Without proper controls or orthogonal validation, these false positives cannot be distinguished from true interactions. Thus, characterizing and addressing these technical challenges and designing proper controls for future investigations of PPI dynamics across different biological contexts is highly justified.

Here, we demonstrate the importance of correcting for changes in the background proteome within AP-MS experiments and provide solutions integrating statistical and experimental strategies. As a case study, we focused on the sole intracellular O-GlcNAc transferase, OGT. Beyond OGT’s catalytic role adding a single N-acetylglucosamine (GlcNAc) to thousands of nuclear and cytoplasmic proteins, it also interacts with hundreds of proteins (17) which may inform its important non-catalytic functions (18). Because tight post-transcriptional regulation of *Ogt* counteracts transgene overexpression and proper localization (19–21), we engineered a dual epitope tag enrichment-control system. This strategy allows for investigation of OGT PPIs at equivalent, endogenous protein levels, and the use of a control which accurately reflects the affinity matrix-specific bead background. We show that acute inhibition of OGT via OSMI-4, a selective small molecule inhibitor that induces major changes to cell biology (22,23), leads to changes in global protein abundance and, thus, the bead background. These differences were accounted for through parallel proteome profiling and/or by applying traditional AP-MS controls on a per-condition basis, both of which aided results interpretation and explained previously unidentified sources of biological and technical variation. Finally, we take a feasible statistical modeling approach that facilitates a direct comparison between biological states of interest, allowing for the confident identification of context-dependent OGT PPIs.

## Experimental Procedures

### Mouse embryonic stem cell culture

ES-E14TG2a mouse embryonic stem cells were cultured feeder-free on plates coated in 0.2% porcine gelatine in “serum/LIF” conditions: knockout DMEM medium (Gibco, 10829-018), 10% HyClone characterized heat-inactivated fetal bovine serum (FBS, Cytiva, SH30396.03), 1x GlutaMAX (Gibco, 35050-061), 1x MEM non-essential amino acids (NEAA, Gibco, 11140-050), 0.1mM beta-mercaptoethanol (Gibco, 21985-023), and leukemia inhibitory factor (LIF, made in house). Cells were maintained in a 37°C, 5% CO_2_ incubator. Low passage numbers were maintained, and cells were routinely checked for mycoplasma contamination.

### Generation of affinity tagged-OGT mESC lines

The pDonor-tBFP-NLS-Neo plasmid was provided by the Nakayama lab (Addgene, 80766). The plasmid was PCR amplified to generate overhangs that are complementary to one of two sequences: one, homology arms amplified from the endogenous *Ogt* locus, and two, a gBlock (Integrated DNA Technologies) containing a puromycin resistance gene, a T2A sequence, and either a 3x-FLAG or 2x-Strep tag (**Supplementary Figure 1** for primers and sequences). The repair template was then cloned into the pDonor plasmid via Gibson assembly (New England Biolabs (NEB)). Oligos harboring the sgRNA sequences (IVT) were flanked by overhangs for in-vitro transcription via the EnGen sgRNA synthesis kit (NEB, E3322V). Four microliters of IVT-synthesized sgRNA (200 ng/µl) were complexed with 1 µl of 40 µM recombinant Cas9 (UC Berkeley QB3) for 15 minutes at 37°C. The resulting ribonucleoprotein (RNP) was then immediately added to the nucleofection solution.

Wild-type (WT) E14 mESCs were cultured on 10 cm^2^ plates until 80% confluency. To generate a single-cell suspension, cells were trypsinized (Gibco, 25200114) and resuspended in fresh media. Cells were counted via an automatic cell counting system (EVE, NanoEntek), and aliquots of 1e5 cells were washed once with 37°C PBS (Gibco, 10010023) and then resuspended in Lonza Nucleofector Solution + Supplement from the P3 Primary Cell 4D-Nucleofector™ X Kit (Lonza Bioscience, V4XP-3032). The solutions mixed with cells to be edited also contained the RNP complex and 3 µl (250 ng/µl) repair template. The negative control for antibiotic selection contained all components besides the repair template, and the positive control for nucleofection efficiency contained pmaxGFP plasmid (1 µg) instead of the repair template. Upon resuspending cells in the Nucleofector Solution, cells were transferred to the 16-well Nucleocuvette and nucleofected using the Amaxa 4D-Nucleofector (Lonza Bioscience), P3 program. Cells were then immediately resuspended in 80 µl 37°C media and allowed to rest in the incubator for ten minutes. Cells were then plated on gelatinized 12-well plates for a two-day recovery period.

For antibiotic selection, cells were transferred to a 10 cm^2^ plate, and, after 18 hours, were selected for with 1 µg/mL Puromycin for seven days. The media was changed every 48 hours and cell death in negative controls was monitored daily. Individual colonies on the plate were considered to be derived from single cells, and colonies were transferred one-by-one to gelatinized 96-well plates for expansion and validation. Once single cell clones were confirmed to have had the correct tag insertion, at least three clones were mixed together to avoid clonal effects.

### PCR and western blot analysis of genome editing

To check whether the affinity tag construct had been knocked into the endogenous *Ogt* locus, 5e4 cells were lysed in 50 mM Tris pH 8, 0.1% SDS, 1 µl/mL Proteinase K (NEB, P8107S) and incubated at 37°C for 1 hour, then 80°C for 20 minutes. Lysate was briefly vortexed and then 1 µl was added to a PCR reaction mixture containing primers that anneal outside of the repair template homology arms **(Supplementary Figure 1B)** and Q5 High-Fidelity Master Mix (NEB, M0492S). The cycling conditions were as follows: one cycle at 98°C for 30 seconds; 34 cycles of 98°C for 10 seconds and then 72°C for 90 seconds; 72°C for 2 minutes, then hold at 4°C. PCR amplicons were separated on a 2% agarose gel containing ethidium bromide, and amplicon sizes were estimated using a 1 kb+ ladder (NEB, N0469S). Since *Ogt* is X-linked and the mESCs are male, there is no possibility of heterozygous editing. Editing was confirmed by an amplicon molecular weight shift above wild type and subsequent Sanger sequencing.

Cells which had the affinity tag construct knocked into the endogenous *Ogt* locus were then checked for tag co-expression with OGT. 1e6 cells were lysed with RIPA buffer supplemented with Halt Protease and Phosphatase Inhibitor Cocktail (Thermo Fisher Scientific, 78446) and 1:1000 benzonase (Sigma Aldrich, E1014), sonicated using the PIXUL multi-sample sonicator (Active Motif) for two minutes at 14°C (Pulse [N]: 50; PRF [kHz]: 1.00; Burst Rate [Hz]: 20.00), and then centrifuged at maximum speed for 15 minutes at 4°C. Protein concentration was quantified via BCA assay (Thermo Fisher Scientific, 23227) and 20 µg protein was boiled in 1x LDS sample buffer (Thermo Fisher Scientific, NP0007) + 3.75% β-ME and subsequently loaded onto a 4-12% NuPAGE Bis-Tris gel for SDS-PAGE. Protein was transferred to a nitrocellulose membrane by semi-dry transfer (Thermo iBlot2, IB21001) and blocked with 5% BSA, 0.1% NaN_3_ in 1x TBS for 30 minutes at room temperature with shaking. The membrane was incubated with rabbit anti-OGT (Cell Signaling Technologies, 24083S, 1:3000), mouse anti-TUBA1B (alpha-tubulin) (Proteintech, 11224-1-AP, 1:20,000), and either mouse anti-FLAG M2 (Sigma Aldrich, F3165, 1:3000) or mouse StrepMAB (IBA, 2-1507-001, 1:3000) at 4°C overnight with shaking. The membrane was washed 3x with TBS-T (1x TBS + 0.1% Tween-20) and then incubated with Goat anti-Rabbit 800 (LI-COR, 926-32211) and Goat anti-Mouse 680 (LI-COR, 926-68070) secondary antibodies for 45 minutes at room temperature with shaking. The membrane was again washed 3x with TBS-T and then 1x with TBS before imaging on a LI-COR Odyssey Fc system.

### Cell lysis and affinity purification

Cells were cultured in 15 cm^2^ plates prior to harvest. Twenty-four hours before harvest, 20 µM OSMI-4 (MedChemExpress, HY-114361) or the equivalent volume of DMSO (0.1% DMSO of total culture volume) was added directly to the culture medium. Triplicates of each sample group were harvested by washing adherent cells with ice-cold PBS, then scraping in ice-cold PBS and centrifuging at 400 *g* for five minutes at 4°C. Cells were transferred to a 1.5 mL Protein Lo-Bind Eppendorf tube and centrifuged at 500 *g* for three minutes at 4°C, followed by aspirating off all PBS. Cell pellets were then resuspended in 5x packed-cell-volume of modified RIPA buffer (no SDS) supplemented with 1x Halt Protease and Phosphatase Inhibitor Cocktail, 20 µM OSMI-4, and 20 µM PUGNAc (Sigma Aldrich, A7229) and incubated on ice for 15 minutes. Samples were transferred to a round-bottom 96-well plate and then sonicated in a PIXUL multi-sample sonicator for five minutes at 14°C. Samples were transferred back to Eppendorf tubes and 1:250 benzonase and 10 mM MgCl_2_ were added. Samples were incubated on an end-over-end rotator for 45 minutes at 4°C and then were centrifuged at maximum speed at 4°C for 20 minutes. Protein quantification was done via BCA assay, and each sample was normalized to 2 mg in 500 µl lysis buffer. At this point, 20 µg of lysate from the FLAG-OSMI-4 and FLAG-DMSO samples were reserved as “proteome-level samples” and were processed via protein aggregation capture (see Methods: Proteome sample preparation). Twenty (20) µl of anti-FLAG magnetic bead slurry (Sigma Aldrich, M8823) or 20 µl Protein A/G magnetic bead slurry bound with mouse isotype antibody (Sigma Aldrich, SAB4702114) was equilibrated in lysis buffer twice before adding their respective samples. Affinity purification was carried out overnight at 4°C with end-over-end rotation. The next day, the beads were briefly spun down in a tabletop microcentrifuge and then placed on a magnetic rack. The flow-through was retained for analysis. Beads were washed 2x with 200 µl of Wash Buffer I (50 mM Tris pH 8, 150 mM NaCl, 1 mM EDTA, 0.1% Triton X-100), then 2x with 200 µl of Wash Buffer II (50 mM Tris pH 8, 150 mM NaCl in HPLC-grade H_2_O). Before the last wash, beads were transferred to new tubes. 5% bead slurry was reserved for western blot analysis (O-GlcNAc MultiMab: Cell Signaling Technologies 82332S 1:1000; OGA: Cell Signaling Technologies 60406S 1:1000) and the remainder of the beads were suspended in 50 µl Wash Buffer II and stored at -80°C until further processing.

For proteome samples comparing the 3x-FLAG–OGT and 2x-Strep–OGT mESC lines, quadruplicate samples were cultured in 10 cm^2^ plates and were not treated with DMSO or OSMI-4. Lysis was carried out in the exact same manner as the AP samples, except for the RIPA buffer contained 0.1% SDS. Forty micrograms of lysate was reserved for protein aggregation capture (PAC) (29).

### Generation of HCF-1-V5 mESC line and reciprocal immunoprecipitation

Construction of the HCF-1-V5, 3x-FLAG-OGT mESC cell line was accomplished using the CRISPaint system (24), which utilizes non-homologous end joining for rapid gene tagging. Briefly, a sgRNA targeting the stop codon of *Hcfc1* (TCTAAGGCTGATGGTCAGTG) was cloned into the target selector plasmid via ligation-independent cloning (25). A gBlock containing the sequences of a V5 tag and Blasticidin resistance gene tethered by a T2A self-cleaving peptide were cloned into a universal donor plasmid via Gibson assembly. These plasmids, alongside the +1 frame selector plasmid, were transfected into 3x-FLAG-OGT mESC using Lipofectamine 2000. Two days later, edited cells were selected for by a 5-day 5 µg/ml blasticidin treatment. Cells were single cell cloned and validated for correct tag insertion into the *Hcfc1* locus via PCR and western blot as described above. Ten clones were mixed together to avoid clonal effects.

For validation of the OGT-HCF-1 interaction with or without OSMI-4 treatment, reciprocal anti-V5 immunoprecipitation was performed. HCF-1-V5, 3x-FLAG-OGT mESC or 3x-FLAG-OGT mESC (control), DMSO or 20 µM OSMI-4 treatment samples were prepared in the same manner as described above. 500 µg of cell lysate was normalized to 400 µl volume and then added to 50 µl (slurry) pre-equilibrated anti-V5 magnetic beads (MBL Life Science, M215-11). The reaction mixture incubated overnight at 4°C, after which the flow-through was removed and beads were washed twice with Wash Buffer I and once with Wash Buffer II. Protein was eluted by resuspending beads in 50 µl of SDS-PAGE loading buffer (Thermo Fisher Scientific, NP0007, plus 3.75% β-ME) and boiling at 95°C for 5 minutes with shaking. Inputs and 25% of the bead boil were blotted with anti-V5 (Cell Signaling Technology 13202S, 1:1000) antibody (HCF-1), anti-FLAG M2 (Sigma Aldrich, F3165, 1:3000) antibody (OGT), and anti-Alpha Tubulin (Proteintech, 11224-1-AP, 1:20,000) antibody as a loading control. The blot analyzing HCF-1 cleavage patterns utilized an anti-HCF-1 antibody raised against the HCF-1 C-terminal half (Bethyl Laboratories, A301-399A, 1:2000). Densitometry analysis was performed using ImageJ (26). Blots are representative of two independent experiments.

### AP-MS sample preparation

AP beads containing the co-immunoprecipitated proteins were thawed on ice and 50 µl of 8 M urea in 50 mM Tris pH 8 was added to each sample. Lys-C (1 mAU) (Fujifilm, 121-05063) was then added to each sample and incubated at 37°C for 30 minutes with shaking. The samples were placed on a magnetic rack, and the supernatant was moved to a new tube, continuing the digestion at 37°C with shaking. The beads were resuspended in 50 µl 50 mM Tris pH 8 containing 500 ng trypsin (Promega, V5113) and incubated at 37°C for 30 minutes with shaking. The supernatant was then added to the previously digesting solution, and the beads were washed once with 50 µl 50 mM Tris pH 8, which was also added to the digestion mixture. The digestion continued overnight at 37°C with shaking. The next day, peptides were reduced and alkylated by addition of 10 mM TCEP and 10 mM iodoacetamide and incubation at room temperature in the dark while shaking. Digestion was quenched by adding 10% trifluoroacetic acid (TFA) to a final concentration of 1%, and the solutions were centrifuged at 18,000 *g*, 4°C for 10 minutes prior to desalting. Peptide desalting was performed using StageTips made in-house (27,28); in brief, StageTips were conditioned with methanol, 50% ACN + 0.1% TFA, and then 0.1% TFA prior to sample loading. Samples were washed twice with 0.1% FA before eluting in 100 µl 50% ACN + 0.1% FA. Peptides were then frozen, dried via vacuum centrifugation and resuspended in 10 µl 0.1% formic acid (FA). Peptides were quantified via Nanodrop and normalized across samples such that there was ∼500 ng peptide per 4 µl injection volume.

### MS-based proteomics sample preparation

For proteome samples comparing the 3x-FLAG–OGT and 2x-Strep–OGT mESC lines, 40 µg of lysate in RIPA buffer containing 0.1% SDS was used for protein aggregation capture (PAC) (29). Samples were reduced with 5 mM dithiothreitol (DTT) for 30 minutes at 56°C, cooled on ice, and briefly centrifuged prior to alkylation with 15 mM iodoacetamide (IAA) for 20 minutes at room temperature in the dark. Residual IAA was quenched with DTT for 15 minutes at room temperature in the dark. Prior to PAC, the buffer pH was titrated to approximately 9.0 with 1M sodium hydroxide (NaOH). PAC was performed using hydroxyl terminated magnetic beads (MagResyn, MR-HYX005) at a protein-to-bead ratio of 1:4 (w/w). Beads were equilibrated in 70% acetonitrile (ACN), and protein lysate, which was adjusted to a final concentration of 70% ACN, was then added to the beads. Samples were then incubated for 45 minutes at room temperature with continuous agitation on a thermomixer at 550 rpm. Protein-bead aggregates were first washed with 100% ACN and then 70% ethanol. For on-bead digestion, protein-bead aggregates were resuspended in 50 mM HEPES (pH 8.5). Samples were digested with 1 mAU Lys-C (Fujifilm, 121-05063) for 1 hour at 37°C with shaking at 750 rpm, followed by overnight digestion with 300 ng trypsin (Promega, V5113) at 37°C with shaking at 750 rpm. Digestion was quenched by acidification to 1% trifluoroacetic acid (TFA), and beads were removed from the peptide solution via magnetic separation. Samples were centrifuged at 18,000 *g* for 10 minutes at 4°C before peptide desalting.

For proteome-level analysis comparing the DMSO vs OSMI-4 treated 3x-FLAG–OGT mESC samples, 20 µg of the AP input was reserved and concentrated by trichloroacetic acid precipitation. Precipitated protein was resuspended in 100 µl of 8 M Urea, 50 mM Tris (pH 8) with 1 mAU Lys-C (Fujifilm, 121-05063) and digested at 37°C for one hour at 1000 rpm. Immediately following, 300 ng trypsin in 50 µl 50 mM Tris (pH 8) was added to each sample and digestion continued overnight (∼18 hours). Reduction and alkylation were performed by adding 10 mM TCEP and 10 mM IAA to each sample and incubating for 30 minutes at room temperature in the dark while shaking. Peptides were acidified to 1% TFA and centrifuged at 18,000 *g* for 10 minutes at 4°C prior to desalting.

Peptide desalting was performed as described above. Eluted peptide solutions were frozen, dried by vacuum centrifugation, and then resuspended in 0.1% FA prior to peptide quantification using the Pierce Quantitative Colorimetric Peptide Assay (Thermo Fisher Scientific, 23275). Peptide concentrations were adjusted to 200 ng/µl prior to LC-MS/MS analysis on the Orbitrap Astral.

### LC-MS/MS

#### Data-dependent acquisition on Orbitrap Eclipse

AP-MS samples were analyzed by LC-MS/MS on an Orbitrap Eclipse Tribrid-mass spectrometer (Thermo Fisher Scientific) coupled online to an Easy nLC 1200 (Thermo Fisher Scientific). Samples (4 µl) were injected onto an Aurora Ultimate 25x75 XT C18 column (IonOpticks, AUR4-25075C18-XT) heated to 60°C. Peptides were progressively eluted over 110 minutes using a gradient containing HPLC solvent A (0.1% FA) and HPLC solvent B (80% ACN, 0.1% FA). The gradient structure was as follows: 6% solvent B at 1 minute (80% ACN, 0.1% FA) to 36% B at 85 minutes, then 72% B until minute 94, 90% B until minute 100, and finishing at 60% B. For mass spectrometry, a precursor scan (scan range: *m/z* 350-1800) was first performed in the Orbitrap at resolution of 60,000, with an AGC target of 4e5. For precursor selection, monoisotopic peak determination was set to “peptide” with the most abundant peak used for the isolation window center. Analytes had to meet a minimal intensity of 5e4, be within a charge state range of 2-8, and dynamic exclusion was enabled with a 30s duration. Precursors were sorted by highest charge state and lowest m/z for subsequent MS/MS analysis. Precursors underwent high-collision dissociation (HCD) fragmentation with a normalized collision energy of 28, and product ions were detected measured in the Orbitrap at a resolution of 15,000. The quadrupole precursor isolation window was set to 1.7 *m/z*, and the maximum MS2 injection time was 105 ms, targeting an AGC of 1e5. Electron-transfer/Higher-energy collision dissociation (EThcD) was also performed, triggered by detection of the HexNAc oxonium ion (204.0867 *m/z*) or HexNAc fragment (138.0545 *m/z*) (15 ppm tolerance) as previously described (30). However, information from these scans are not presented in this report.

#### Data-independent acquisition on Orbitrap Astral

Proteome samples were analyzed by LC-MS/MS on an Orbitrap Astral mass spectrometer (Thermo Fisher Scientific) coupled online to a Vanquish Neo UHPLC system (Thermo Fisher Scientific). Samples (200 ng) were injected onto an Aurora Ultimate 25x75 XT C18 column (IonOpticks, AUR4-25075C18-XT) heated to 60°C. HPLC solvents A (0.1% FA) and B (80% ACN, 0.1% FA) were used to construct a 72-minute total gradient with the following structure: 4% B was applied for 1.7 minutes, then 8% B until minute 12.5, 25% B until minute 52.7, and 35% B until minute 64.7. The column was then washed with 99% B until gradient end at 71.1 minutes. Mass spectrometry was performed with Lock Mass correction on, via the internal EASY-IC source. A MS survey scan was performed in the Orbitrap (scan range: *m/z* 380-980) at a resolution of 120,000, with a normalized AGC target of 500% (absolute: 5e6) and a maximum injection time of 5 ms. Narrow window data-independent acquisition (nDIA) was performed in the Astral mass analyzer within a precursor mass range 380-980 *m/z*, with an isolation window of 2 *m/z* and a window overlap of 0.5 *m/z*. HCD normalized collision energy was set to 25% and the normalized AGC target was 1000% (absolute: 1e5). The maximum injection time was 3.5 ms and loop control was set to “time” with a boundary of 0.6 seconds.

### Data analysis

#### MS data searching

AP-MS data analyzed on the Eclipse were searched in MaxQuant v2.6.5.0 (31) against the mouse UniProt database (12/28/2017) with 264 common laboratory contaminants and 553 smORFs added. Digestion specificity was set to “Trypsin” and “Lys-C” with two missed cleavages allowed. Carbamidomethylation (C) was added as a fixed modification and Oxidation (M) and Acetylation (protein N-term) were added as variable modifications, with a maximum number of 5 modifications per peptide. Label-free quantification (LFQ) was enabled with a minimum ratio count of two, classic normalization, and Fast LFQ (32). The MS1 and MS2 mass tolerances were set to 20 ppm, and the protein and peptide false discovery rate was set to 1%. The match-between-runs feature was enabled with default settings. The proteinGroups file containing protein-level LFQ-normalized intensities was then used for all downstream analyses.

Proteomics data (both the 3x-FLAG-OGT vs. 2x-Strep-OGT cell lines and the 3x-FLAG-OGT OSMI-4 vs. DMSO experiments) analyzed on the Astral were searched using CHIMERYS v4.0 on the Ardia Server in Proteome Discoverer v3.2.0 (Thermo Fisher Scientific). Data were searched against the *Mus musculus* Swiss-Prot reference database (downloaded 02/05/2025, 17,207 entries) using the Inferys v4.7.0 prediction model. Enzyme specificity was set to “Trypsin/P” with two maximum missed cleavages. Carbamidomethylation (C) was set as a static modification and Oxidation (M) as a variable modification, with maximum three variable modifications per peptide. The fragment mass tolerance was set at 15 ppm, and the target FDR for peptides, PSMs, and proteins was 1%. All peptides were used for protein roll-up, and the average of the top 5 most abundant peptides were used for protein-level quantification. Identifications were checked against a database of 264 laboratory contaminants. The strict parsimony principle was set to TRUE for protein grouping. The “Proteins” file containing raw protein-level intensity values per sample was used for downstream analysis. The only differences in the search parameters of the two experiments was that the 3x-FLAG-OGT vs. 2x-Strep-OGT cell line experiment set a maximum fragment mass tolerance of 20 ppm, and MS2-level area-based quantification was performed across all files (“Quan in all files”).

#### Experimental Design and Statistical Rationale

For AP-MS and MS-based proteomics experiments, three independent biological replicates were analyzed per condition: 3x-FLAG-OGT and 2x-Strep-OGT mESC lines, with DMSO control or OSMI-4 treatment. A fourth replicate was included for the comparison of the 3x-FLAG-OGT and 2x-Strep-OGT mESC lines to ensure comparison of these highly similar mESC lines would not be underpowered. Detailed descriptions of controls and efforts taken to reduce technical variation are provided throughout the Results section.

All data analysis was done in RStudio - R v4.5.2. For AP-MS Eclipse samples, proteins which were identified as laboratory contaminants or their peptides identified by reverse sequences were filtered out, as well as any proteins that did not have at least two non-zero values in at least one condition (a condition being Enrichment + Treatment + Beads, for example: anti-FLAG AP, DMSO treatment, FLAG beads). Quantification values were normalized via variance-stabilizing normalization (VSN) and then log_2_ transformed. Data were imputed by first identifying the most appropriate imputation method per data feature (missing-not-at-random/MNAR or missing-at-random/MAR) using the model.Selector feature in the ImputeLCMD R package (33,34). MAR data was imputed via k-nearest neighbors, and MNAR data was imputed via quantile regression imputation of left-censored data (QRILC). The after-imputation data distributions remained Gaussian and with a nearly identical dynamic range as to the before-imputation data. For proteome Astral samples, proteins which were identified as contaminants were filtered out, as well as any protein that did not have at least two non-zero values in at least one condition. Data were log_2_ transformed and then median normalized (non-zero method). For all samples, statistical analyses to generate pairwise comparisons were performed via *limma* (35). Moderation via eBayes was performed with trend = TRUE. Data were corrected for multiple hypothesis testing by the Benjamini-Hochberg method. The results from each contrast were combined into a single data frame by joining on a protein’s UniProt accession number. To generate a database of known OGT interactors, which we termed “Reported Interactors”, protein interactors associated with OGT were downloaded from the BioGRID (downloaded April 14th, 2026, mouse and human) (36) and IntAct (downloaded April 14th, 2026, mouse and human) (37) databases and integrated with interactions detected in our previous study (38). Interactions were filtered to include only those identified through physical interaction detection methods (*i.e.*, excluding methods such as proximity labeling, functional associations, subcellular co-localization, *etc.*). Gene symbols for interactors detected with human OGT were converted to the corresponding mouse gene symbol and de-formatted (i.e. capitalizations) for integration with the mouse datasets. The final dataset was de-duplicated such that one protein is not counted as an interactor more than once. All figures were generated using ggPlot2 (39) or GraphPad Prism v10.6.0. Figure aesthetics were modified in Adobe Illustrator (2025).

To facilitate reproducible statistical analyses involving similar linear model structures, we developed an R shiny application (https://github.com/melinabrunelli/LinearModelR) that runs on any feature-by-sample data matrix with accompanying metadata. In addition to comma- and tab-delimited files, the application accepts input in Gene Cluster Text (GCT) format. Users can define main and secondary effects, which form the basis of an interaction-term linear model, and can specify contrasts in a delta-delta format. The application also supports repeated-measures designs (e.g., repeated time point or inter-condition samples from the same patient), although this feature was not used in the present work.

## Results

### CRISPR-Cas9-mediated knock-in of orthogonal affinity epitopes into the endogenous Ogt locus in mouse embryonic stem cells

The overexpression of an epitope-tagged protein for PPI characterization can induce unintended protein abundance changes, which can lead to the identification of differential background binders as interactors. To avoid this and other challenges associated with protein overexpression, and to circumvent the tight post-transcriptional control of OGT levels (21,40), we employed the CRISPR/Cas9-mediated homology directed repair (HDR) system to install epitope tags into the endogenous *Ogt* locus **(Figure 1A)**. We tagged OGT on the N-terminus because it is farther away from the catalytic domain, as this may have less of a chance of interfering with its catalytic function. Additionally, the N-terminus is slightly less conserved than the C-terminus, and previous reports demonstrated successful tagging of OGT on the N-terminus with no apparent changes to its known interactome (41). We generated two epitope-tagged OGT cell lines—3x-FLAG-OGT and 2x-Strep-OGT—so that one epitope/line is used to perform the affinity enrichment and the other can act as a background control, as it is incubated with the same affinity matrix as the enrichment, but OGT cannot be enriched because it harbors a different epitope. We electroporated mouse embryonic stem cells (mESC) with a ribonucleoprotein (RNP) complex consisting of an *in vitro*-transcribed sgRNA—which targets the start codon of *Ogt*—and recombinant Cas9 protein (42), as well as a HDR repair template. The repair template contained homology arms that recognize the region immediately surrounding the *Ogt* start codon, and the homology arms flanked a puromycin resistance gene fused to either the 3x-FLAG or 2x-Strep tag by a T2A self-cleaving peptide **(Figure 1B, Supplementary Figure S1A)**, allowing for the insertion of the affinity epitope and selection marker directly following the start codon. We confirmed successful integration of each epitope tag by PCR **(Figure 1C, Supplementary Figure S1B)** and western blot analysis **(Figure 1D)**. Additionally, we performed narrow window data-independent acquisition (nDIA) mass spectrometry on the 3x-FLAG-OGT and 2x-Strep-OGT mESC lines and found that there are very few differences between their proteomes **(Figure 1E),** permitting us to utilize these lines as an enrichment/control pair.

**Figure 1.**
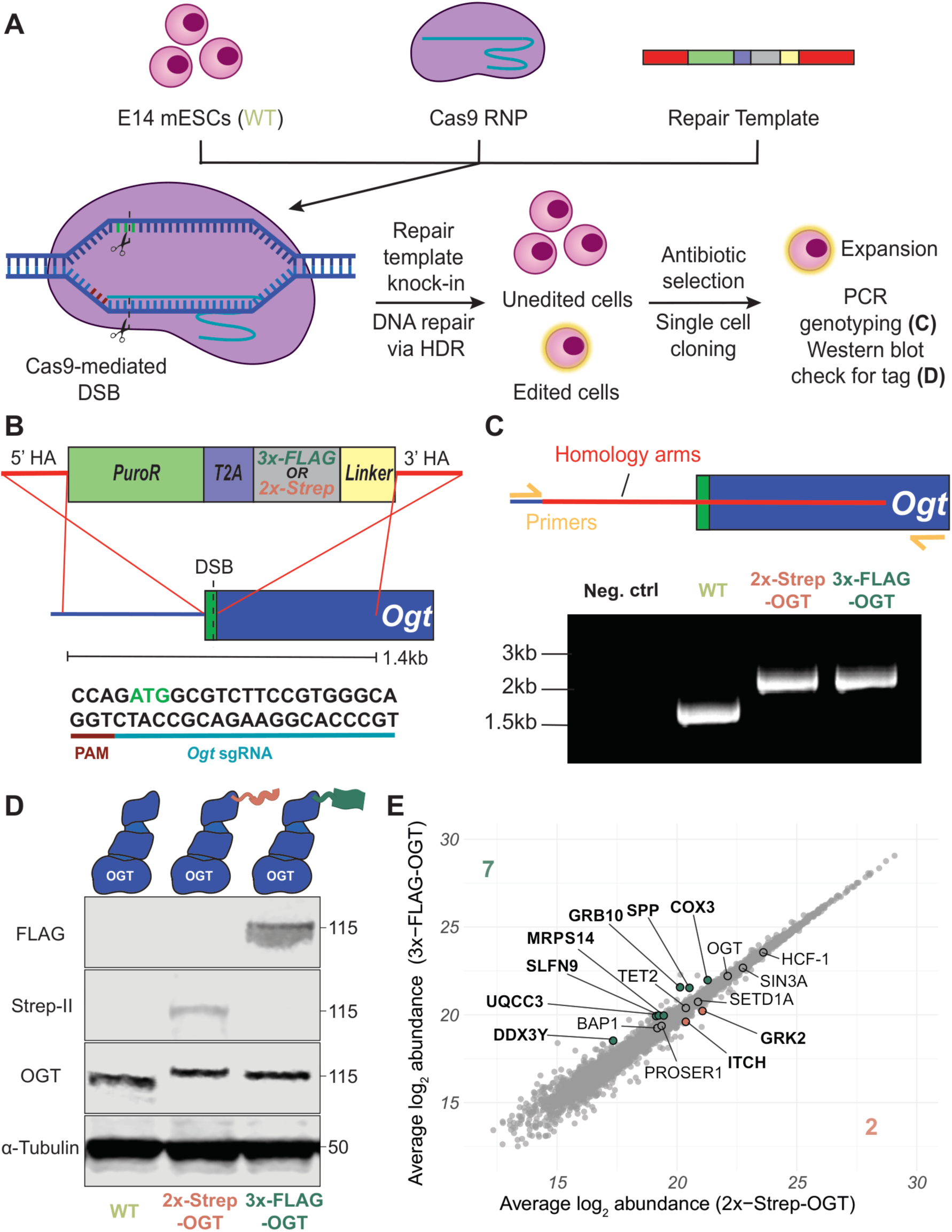
CRISPR-Cas9-mediated knock-in of orthogonal affinity epitopes into the endogenous *Ogt* locus in mouse embryonic stem cells. **A. Experimental scheme depicting the CRISPR-Cas9-mediated homology directed repair (HDR) workflow.** An *in-vitro* transcribed sgRNA is complexed with recombinant spCas9 to form a Cas9 ribonucleoprotein (RNP), which, alongside the repair template, is nucleofected into mouse embryonic stem cells (mESCs). A double-stranded break (DSB) is introduced near the *Ogt* start codon and the repair template is inserted via HDR repair pathway. Cells with genomic integration of the repair template are selected for with antibiotics, and single cells are then genotyped via PCR and checked for OGT-tag co-expression via western blot. **B. Schematic depicting the HDR repair template, its destined location in the *Ogt* locus, and the sgRNA sequence targeting the *Ogt* start codon.** The repair template consists of 5’ and 3’ homology arms (HAs), a puromycin resistance gene (PuroR), a T2A self-cleaving peptide, either a 3x-FLAG or a 2x-Strep affinity epitope sequence, and a short, flexible linker. **C. PCR amplification of mESC genomic DNA for tag knock-in validation.** Primers that anneal directly outside of the beginning and end of the repair template region allows for confirmation of affinity tag knock-in by comparing WT (expected size: 1610 bp) to 2x-Strep–OGT (expected size: 2378 bp) or 3x-FLAG–OGT (expected size: 2360 bp) mESC. The negative control included master mix and primers, but no template DNA. **D. Western blot analysis of OGT - affinity tag co-expression.** Whole cell lysate from WT, 3x-FLAG–OGT, or 2x-Strep–OGT mESC was probed by western blot using the indicated antibodies, showing that the affinity tags are co-expressed with OGT and cause a slight molecular weight shift compared to WT OGT. **E. Scatter plot comparing 3x-FLAG-OGT and 2x-Strep-OGT mESC proteomes.** Quadruplicates of each cell line were analyzed by nDIA and the protein normalized average abundance is plotted. Labels in bold indicate proteins with adj. p-value < 0.05. Green points = enriched in 3x-FLAG line; orange points = enriched in 2x-Strep line.

### Proper bead background representation is crucial to distinguish between true and false positives for characterization of OGT PPIs by AP-MS

To exemplify how our endogenous epitope-tagged lines enable an effective AP-MS experimental scheme for OGT PPI identification, we performed AP-MS on whole cell lysates **(Figure 2A)**. OGT and its interactors were enriched from the 3x-FLAG-OGT (“fOGT”, where f = 3x-FLAG) mESC line using anti-FLAG (“FLAG”) antibody-conjugated magnetic beads; this enrichment is hereby referred to as “fOGT_FLAG_”, where the subscript indicates the bead type. We wanted to recapitulate previous findings which showed that the bead background depends on the affinity purification matrix (10,43), so we utilized two enrichment controls. Our primary enrichment control combined lysate from the 2x-Strep-OGT (“stOGT”, where st = 2x-Strep) mESC line with the same or “matched” anti-FLAG antibody-conjugated beads, termed “stOGT_FLAG_”. For demonstration, we also included an insufficient control where 3x-FLAG-OGT mESC lysate was incubated with “mismatched” beads: Protein A/G + antibody isotype (“ISO”) magnetic beads, termed “fOGT_ISO_”. This was to show how differences in the affinity matrices between the enrichment and control can lead to aberrant PPI identifications (44,45). We considered a protein an interactor if it was enriched with fOGT_FLAG_ over the respective control and passed a nominal p-value threshold of 0.05, so that we may monitor all changes in significance across different analyses, as opposed to a more rigorous and appropriate adjusted p-value threshold.

**Figure 2.**
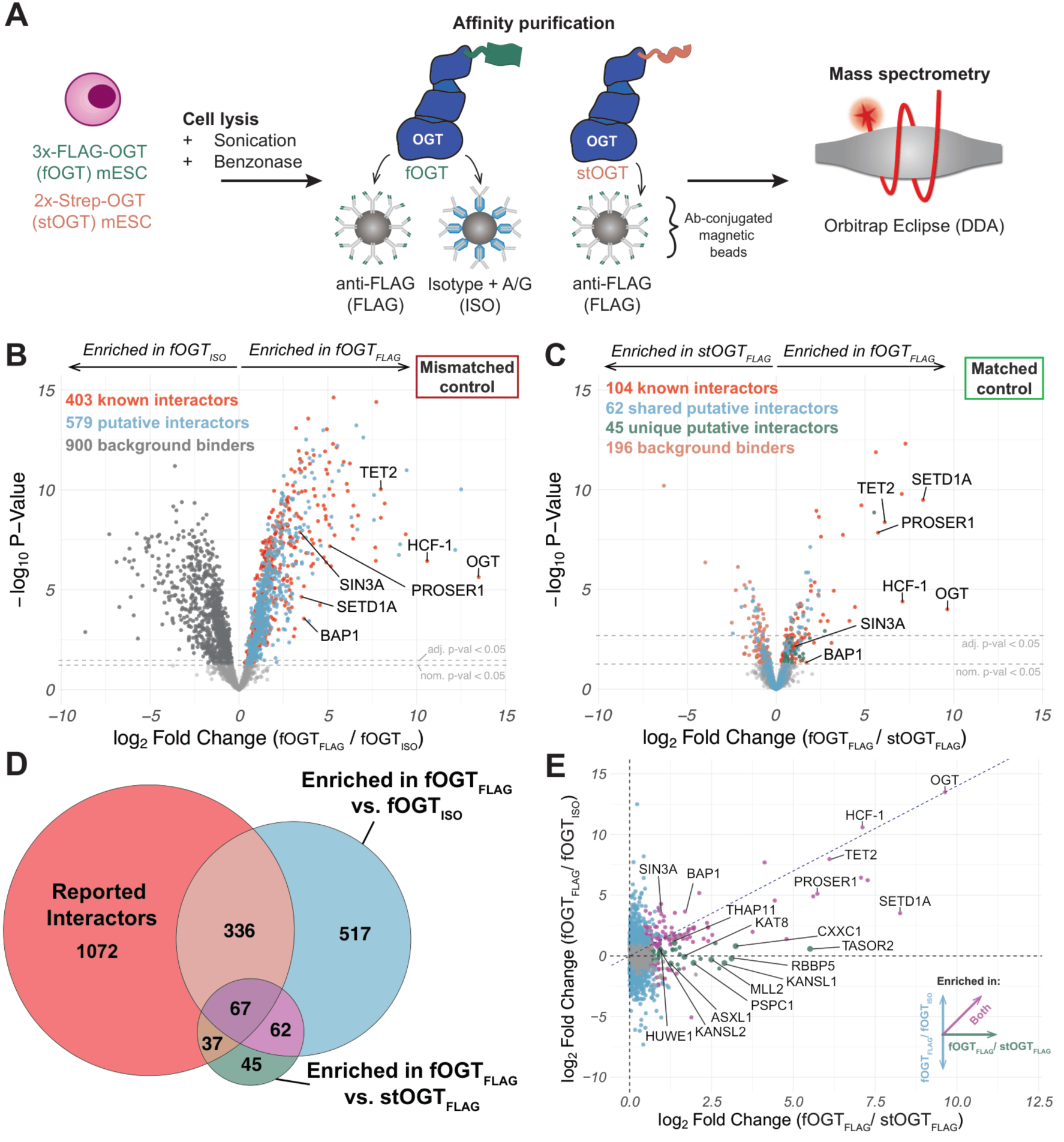
Proper bead background representation is crucial to distinguish between true and false positives for characterization of OGT PPIs by AP-MS. **A. Experimental scheme depicting the AP-MS workflow.** 3x-FLAG–OGT (fOGT) or 2x-Strep–OGT (stOGT) mESC are lysed with RIPA buffer (no SDS) containing benzonase to digest nucleic acids and sonicated to aid nuclear lysis and chromatin shearing. Affinity purification was performed using 3x-FLAG–OGT mESC lysate incubated with anti-FLAG-conjugated magnetic beads (FLAG). The mismatched control combined 3xFLAG–OGT mESC lysate with Antibody isotype + Protein A/G-conjugated magnetic beads (ISO), and the matched control incubated 2x-Strep–OGT mESC lysate with anti-FLAG-conjugated magnetic beads. The samples are then processed for and analyzed by mass spectrometry on an Orbitrap Eclipse. **B. Volcano plot of the fOGT_FLAG_ vs. fOGT_ISO_ pairwise comparison.** Labeled proteins are a subset of reported and validated OGT interactors. Red points are proteins that have been previously reported as an OGT interactor (“known interactor”). Light blue points are enriched proteins that have not been previously reported as an OGT interactor (“putative interactor”). Dark gray points are proteins enriched in the mismatched control (“background binder”). Bolded values specify proteins passing the nominal p-value threshold *p* < 0.05. The top-right corner text box classifies the mismatched control as an improper control. **C. Volcano plot of the fOGT_FLAG_ vs. stOGT_FLAG_ pairwise comparison.** Labeled proteins are a subset of reported and validated OGT interactors. Red points are proteins that have been previously reported as an OGT interactor (“known interactor”). Light blue points are the putative enriched interactors identified in **2B**, 62 of which are also enriched (*p* < 0.05) in this comparison (“shared putative interactor”). Green points are interactors that have not been previously reported and are unique to this comparison (“unique putative interactor”). Orange points are proteins enriched in stOGT_FLAG_ control (“background binders”). Bolded values specify proteins passing the nominal p-value threshold *p* < 0.05. The top-right corner text box classifies the matched control as a proper control. **D. Venn diagram comparing this study’s identified OGT interactors to reported OGT interactors.** Reported interactors (red) consists of interactions reported in the BioGRID and IntAct databases, as well as interactions previously found in a nuclear mESC OGT Co-IP (38) (see **Methods** for additional details). **E. Scatter plot comparing enriched interactors between each enrichment-control comparison.** Y-axis: log_2_ fold changes from the fOGT_FLAG_ vs. fOGT_ISO_ comparison (light blue: log_2_FC > 0 & *p* < 0.05); x-axis: log_2_ fold changes from the fOGT_FLAG_ vs. stOGT_FLAG_ comparison (green: log_2_FC > 0 & *p* < 0.05). Interactors enriched in both comparisons are in dark magenta. The dashed diagonal line intentionally intercepts true zero and OGT to highlight shared interactors. The negative x-axis is hidden for figure clarity.

We first compared fOGT_FLAG_ to the fOGT_ISO_ mismatched control and found a total of 982 proteins enriched with OGT, including several known and validated interactors such as HCF-1, TET2, and SETD1A (**Supplementary Figure S2A**), as well as 900 proteins enriched in the bead background (**Figure 2B)**. We achieved high enrichment efficiency as shown by no observable OGT signal in the enrichment flow-through and presence on the beads, in relation to both controls **(Supplementary Figure S2B)**. To discern which of the 982 proteins may be novel, potentially cell-type specific interactors, we combined reported physical interactions from the BioGRID and IntAct databases (36,46) with OGT nuclear protein interactions from mESC that we described in a recent study (38), which together we called “Reported Interactors”. By comparing proteins enriched in fOGT_FLAG_ to the Reported Interactors dataset, we found that 50% (579/982) of the enriched proteins were not previously reported, or “putative”, interactors. As expected, this large number of novel interactors identified–and the thousands of differential interactors overall–was likely due to the two different affinity matrices harboring specific and unique bead backgrounds. This renders one incapable of discerning true positive interactors from bead-specific background binders (false positives). Indeed, the bead backgrounds of the two controls exhibited very different sets of background binders **(Supplementary Figure S2C)**. Of the 579 putative interactors, 43% (250/579) were not identified or low-abundance in the fOGT_ISO_ samples **(Supplementary Figure S2D)**, and their quantification and thus comparison to fOGT_FLAG_ was only made possible through MaxQuant’s match-by-runs strategy (32) or missing-not-at-random data imputation **(Methods)**. We examined the relative abundance of the 267 remaining putative interactors against all proteins in the 3x-FLAG OGT mESC proteome and observed a trend towards higher abundance **(Supplementary Figure S2E)**. This could suggest that they represent mESC-specific anti-FLAG bead background binders, as protein representation in bead background has a slight bias towards higher abundance proteins (43). Overall, this demonstrates that false positive and false negative PPI identification can occur if one does not utilize a control that represents the affinity matrix-specific bead background.

To further highlight the importance of properly controlling for the bead background, we then compared fOGT_FLAG_ to the matched stOGT_FLAG_ control and saw a nearly 5-fold reduction in the number of differential PPIs identified, with 89% (517/579) of the putative interactors identified in the fOGT_FLAG_ vs. fOGT_ISO_ comparison no longer statistically enriched **(Figure 2C)**. The fOGT_FLAG_ vs. stOGT_FLAG_ comparison yielded 211 OGT protein interactors, 104 of which we previously identified as OGT interactors in mESC nuclei (38). We identified 45 putative interactors unique to the fOGT_FLAG_ vs. stOGT_FLAG_ comparison, most of which localize to the cytoplasm and thus would not have been captured in our previous study. Further comparison of the enriched proteins between the two control comparisons and the Reported Interactors dataset produced 37 previously reported interactors that were identified only in the fOGT_FLAG_ vs. stOGT_FLAG_ comparison **(Figure 2D)**. By plotting the log_2_ fold changes of the fOGT_FLAG_ vs. stOGT_FLAG_ comparison against those of the fOGT_FLAG_ vs. fOGT_ISO_ comparison, we illustrate comparison-specific interactors (data point proximal to the comparison’s respective axis) or interactors identified by both comparisons (data point proximal to the diagonal line intercepting OGT) **(Figure 2E)**. The 37 known interactors–such as subunits of the NSL, COMPASS, and PR-DUB complexes (47–49)–fall along the x-axis and therefore were only identified in the fOGT_FLAG_ vs. stOGT_FLAG_ comparison. Altogether, this indicates that differences between the bead backgrounds can lead to spurious classification of both false positive and false negative PPIs, which can be ameliorated by utilizing a control in which the target protein does not harbor the enrichment epitope but is incubated with the same affinity matrix as the target protein enrichment.

### Global protein abundance changes obfuscate bona fide OGT catalytic inhibition-dependent interactions

We then wanted to demonstrate that the bead background depends not only on the affinity matrix, but also the background proteome, which may differ between the biological contexts in which PPIs are being profiled. For this, we investigated how OGT protein interactions change upon catalytic inhibition of OGT via OSMI-4 (22), a perturbation known to cause significant proteome alterations (21). We treated our fOGT mESCs with 20 µM OSMI-4 or DMSO control for 24 hours, then harvested the cells and performed an OGT enrichment using the same enrichment strategy as previously mentioned. We observed a near complete loss of global O-GlcNAc signal with this OSMI-4 treatment concentration and timepoint **(Figure 3A)**. Relative to the DMSO-treated fOGT_FLAG_ enrichment, the OSMI-4-treated fOGT_FLAG_ enrichment contained more OGT, which was commensurate with its abundance increase, a previously described phenomenon (21,40) **(Supplemental Figure S2B)**. By performing AP-MS and then comparing DMSO- and OSMI-4-treated fOGT_FLAG_, we initially observed significant loss and gain of OGT PPIs, such as an apparent increase in interaction with subunits of histone-modifying epigenetic protein complexes **(Figure 3B)**. In particular, components of the COMPASS and NSL complexes appeared to increase in interaction and more of their complex subunits were detected, as shown by an enrichment of these complexes’ CORUM database terms upon performing gene ontology analysis **(Supplementary Figure S3A)** (50). On the other hand, known OGT interactors TET2, PSPC1, and PROSER1 putatively decreased in interaction with OGT. We detected a potential increase in interaction with proteins involved in cell cycle regulation (CHK1, YES1) and ubiquitin-proteasome activity (FBXO22, UBXN1), two processes for which OGT catalytic activity has been shown to be crucial (51,52). Additionally, components of the Sin3a complex (SIN3A, SINHCAF, SAP30BP, and BRMS1L) and mRNA splicing factors (SRSF6, SRSF9) putatively lost interaction with OGT. We anticipated considerable changes in OGT PPIs because O-GlcNAcylation has been previously implicated in the regulation of protein-protein interactions: promotion or obstruction of those interactions depends on the protein of interest and cellular context (53,54). However, O-GlcNAcylation also regulates protein stability through crosstalk with ubiquitination (55,56) and through inhibiting proteasomal activity (52,57). Thus, we expected that some of these purported interaction changes may actually be due to OSMI-4-induced protein abundance changes altering their representation in the bead background, rather than a true change in interaction.

**Figure 3.**
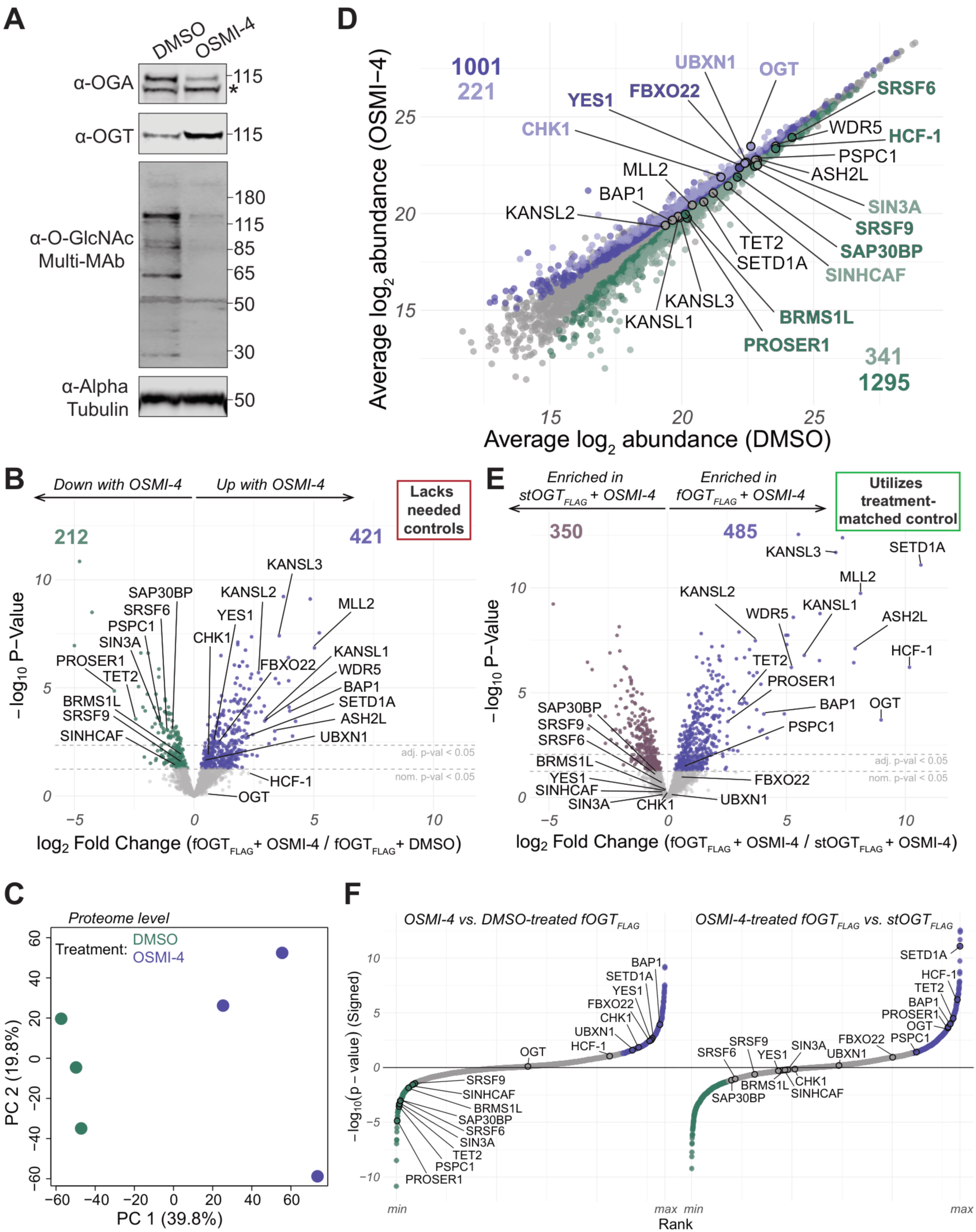
Global protein abundance changes obfuscate *bona fide* OGT catalytic inhibition-dependent interactions. **A. Western blot comparing O-GlcNAc, OGT, and OGA levels between OSMI-4 and DMSO treatment.** Alpha tubulin was used as a loading control; all other antibodies are indicated in figure. The asterisk marks a non-specific band stained by the anti-OGA antibody. **B. Volcano plot of the fOGT_FLAG_ OSMI-4 vs. DMSO treatment comparison.** Interactions gained upon OSMI-4 treatment are in purple; interactors lost upon OSMI-4 treatment are in green. Bolded values specify proteins passing the nominal p-value threshold *p* < 0.05. The top-right corner text box specifies that this comparison lacks needed controls. **C. Principal component analysis (PCA) plot comparing OSMI-4- and DMSO-treated mESC proteomes.** All fully quantified proteins were used for the analysis (8431/8740). Percentages indicate the percent total variance explained by the PC. **D. Scatter plot highlighting proteome-level abundance changes of gained/lost OGT interactors upon OSMI-4 treatment.** The average normalized protein abundance is plotted (OSMI-4 on y-axis; DMSO on x-axis). Bolded labels are proteins whose interaction and abundance significantly change (in the same direction). Light purple/green indicate proteins passing threshold adj. p-value < 0.05; dark purple/green indicate proteins passing nominal p-value < 0.05. Gray points have a nominal p-value of > 0.05. **E. Volcano plot of the OSMI-4-treated fOGT_FLAG_ vs. stOGT_FLAG_ comparison.** Proteins enriched in fOGT_FLAG_ are plotted in purple; proteins enriched in stOGT_FLAG_ (*i.e.*, bead background) are plotted in mauve. Bolded values specify proteins passing the nominal p-value threshold *p* < 0.05. The top-right corner text box specifies that this comparison utilizes a treatment-matched control. **F. Signed log P-value versus rank plot contrasting the fOGT_FLAG_ OSMI-4 vs. DMSO comparison (Left) with the OSMI-4-treatment fOGT_FLAG_ vs. stOGT_FLAG_ comparison (Right).** Signed log P-values were calculated by multiplying the -log_10_*P* by the log_2_ fold change. The two comparisons were ranked separately and plotted against the same y-axis scale. The minimum rank is the protein with the smallest p-value and a negative log_2_ fold change; the maximum rank is the protein with the smallest p-value and a positive log_2_ fold change. Labeled, colored points are the differential interactors/abundant proteins from **3D** and validated OGT interactors whose abundance does not change. Purple = interaction gain with OSMI-4; Green = interaction loss with OSMI-4; Gray = nominal p-value > 0.05.

To confirm this, we reserved some of the AP input to be analyzed as “proteome-level” samples by nDIA on an Orbitrap Astral (7) **(Supplementary Figure S3B)**. Principal component analysis (PCA) verified that the proteomes between the DMSO- and OSMI-4-treated mESC proteomes differed, with PC1 resolving treatment conditions **(Figure 3C)**. Of the hundreds of differentially abundant proteins, 56 proteins with increased abundance and 55 proteins with decreased abundance upon OSMI-4 treatment were also potential gained/depleted interactors identified in the fOGT_FLAG_ enrichments **(Supplementary Figure S3C, S3D)**. Among the seemingly depleted interactors with decreased abundance were PROSER1, the Sin3a complex subunits, and mRNA splicing factors, whereas TET2 and PSPC1 abundances did not change. We also observed increased OGT levels, as well as increased abundance of the putative interactors that regulate cell cycle progression and protein stability **(Figure 3D)**. On the other hand, the histone-modifying complex subunits that appeared to increase in interaction with inhibited OGT did not significantly increase in abundance **(Supplementary Figure S3E)**. Therefore, with parallel proteomic profiling, we were able to identify proteins that appeared to be differentially interacting between two distinct biological contexts but may have actually had altered bead background representation due to protein abundance changes.

Since we determined that the stOGT_FLAG_ control properly accounted for the background proteome in the steady-state OGT enrichment, we hypothesized that treating stOGT_FLAG_ with OSMI-4 would capture the OSMI-4 treatment-specific background proteome and could be used as an optimal control for elimination of false positives. Through comparison of the OSMI-4-treated fOGT_FLAG_ enrichment to the OSMI-4-treated stOGT_FLAG_ control, we properly controlled for the supposed differential interactors whose abundance and interaction changed in tandem **(Figure 3E)**. Compared to the OSMI-4 vs. DMSO fOGT_FLAG_ enrichment, we no longer saw significant enrichment of the putative differential interactors with OGT **(Figure 3F)**. As expected, there were significant differences between OSMI-4 and DMSO-treated stOGT_FLAG_ controls, with the false positive interactors mirroring their enrichment pattern in the fOGT_FLAG_ treatment samples **(Supplementary Figure S3F)**, confirming that the bead backgrounds between the two treatments were indeed different. Proteins such as TET2 and PSPC1, on the other hand, remained enriched with fOGT_FLAG_ even though their interactions with OGT are depleted upon OGT catalytic inhibition. This also held true for PROSER1 despite its decrease in abundance, demonstrating that we could distinguish true changes in interaction with OGT from artifacts arising from different bead backgrounds. In summary, these findings exemplify the need for two important controls when measuring PPIs across changing biological contexts: one, a control for the biological perturbation, such as DMSO for OSMI-4 treatment, and two, a control for the bead background perturbation, such as parallel proteome profiling and/or condition-specific enrichment controls.

### Statistical modeling promotes a background-corrected differential OGT PPI evaluation

Our thorough controls allowed us to distinguish true interactions from background binders in both treatment conditions, but they only enabled a *post hoc* comparison between treatments, *i.e.*, comparing the list of hits from one fOGT_FLAG_ vs. stOGT_FLAG_ treatment condition to the other. We needed a way to directly compare our treatment conditions while still factoring in the biological and technical variation captured by our controls. We considered that this could be accomplished through a linear model analysis: here, we employed an interaction-term linear model, which allowed us to specify a “contrast of contrasts”: within-treatment groups were contrasted (*i.e.*, enrichment vs. enrichment, control vs. control) as well as between-treatment groups (*i.e.*, OSMI-4 vs. DMSO). These contrasts were then further contrasted to produce a direct comparison between fOGT_FLAG_ OSMI-4 vs. DMSO enrichments (**Figure 4A**, **Methods**). As a feasible and user-friendly approach to this modeling strategy, we constructed a Shiny App that employs *limma* (35,58), allowing for complex, multivariate experimental designs (github.com/melinabrunelli/LinearModelR). Upon fitting the data to the model, we observed that the apparent lost interactors (BRMS1L, SINHCAF, SAP30BP, SRSF6, SRSF9) and apparent gained interactors (UBXN1, FBXO22, YES1, CHK1) were no longer enriched in either treatment condition **(Figure 4B)**, in comparison to the non-background-corrected OSMI-4 vs. DMSO-treatment pairwise comparison. Notably, SIN3A and PROSER1 still significantly lost interaction with OGT, illustrating that a distinction can be made between proteins that change interaction because of their altered abundance, and proteins that appear to change interaction because of their differential appearance in the bead background.

**Figure 4.**
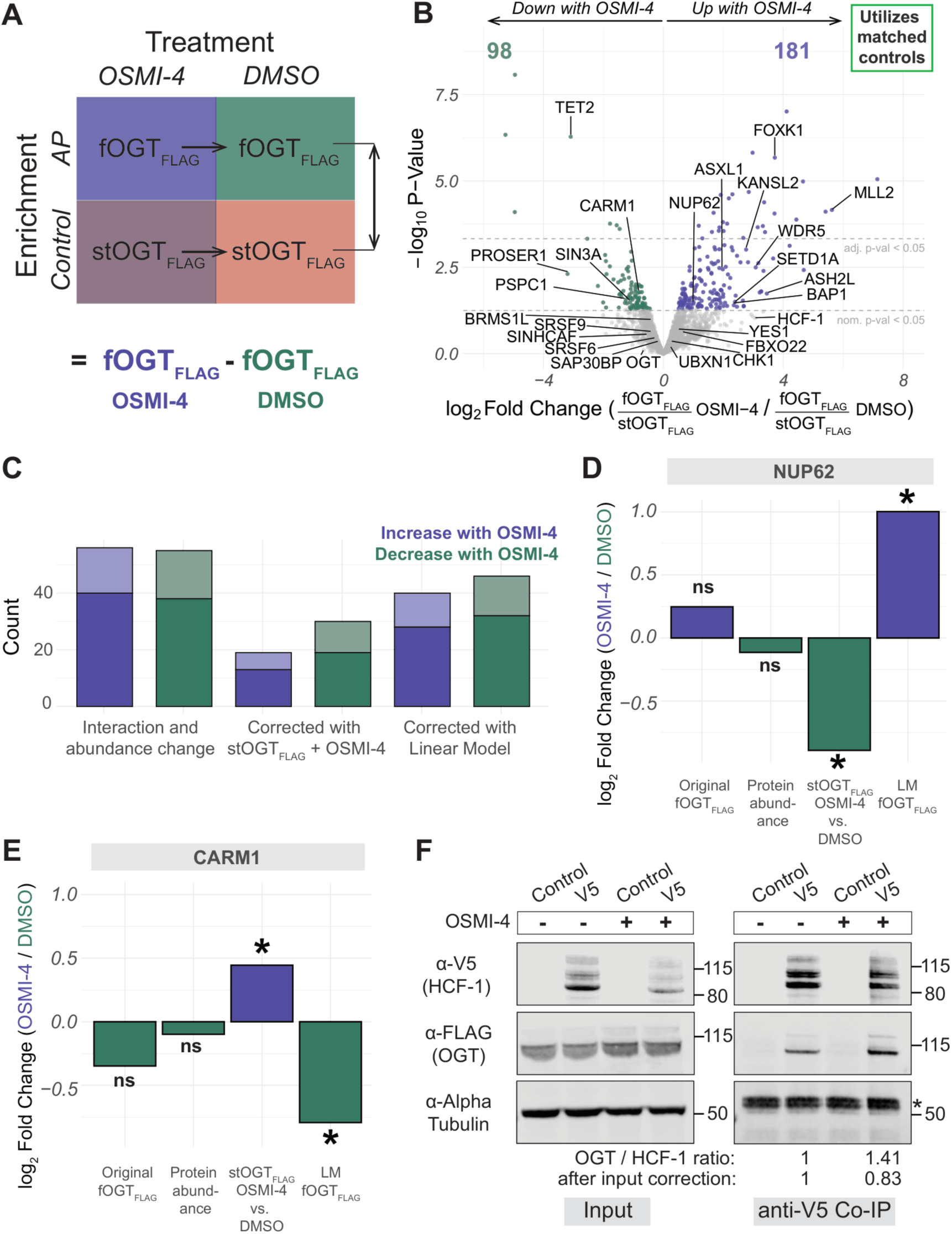
Statistical modeling promotes a background-corrected differential OGT PPI evaluation. **A. Schematic depiction of the linear model strategy.** The statistical model employed in this study models the interaction term between two factors: the enrichment sample (AP or control) and the treatment (DMSO or OSMI-4). Horizontal arrows represent the treatment effect within enrichments, and the vertical double-sided arrow represents the interaction term, also shown as the equation. The model result is the background-corrected contrast between OSMI-4- and DMSO-treated fOGT_FLAG_ enrichments. **B. Volcano plot of the linear model-fitted fOGT_FLAG_ OSMI-4 vs. DMSO comparison.** Interactors gained with OSMI-4 treatment are in purple; interactors lost with OSMI-4 treatment (or more enriched in the DMSO treatment) are in green. Bolded values specify proteins passing the nominal p-value threshold *p* < 0.05. The top-right corner text box indicates that condition-specific controls are used to form this comparison. **C. Bar graph accounting for false positives across the original AP-MS analysis, inclusion of stOGT_FLAG_ controls, and linear model analysis.** Proteins that were originally differential interactors and were no longer statistically significant upon comparison to stOGT_FLAG_ + OSMI-4 or within the modeled AP-MS analysis are considered “corrected”. Light purple/green indicate proteins passing threshold adj. p-value < 0.05; dark purple/green indicate proteins passing nominal p-value < 0.05. Purple = interaction gain with OSMI-4; Green = interaction loss with OSMI-4. **D. NUP62 abundance and OGT interaction status across analyses. “**Original fOGT_FLAG_” = initial fOGT_FLAG_ OSMI-4 vs. DMSO AP-MS analysis (Fig. 3B). “Protein abundance” = average proteome-level abundance (Fig. 3D). “stOGT_FLAG_ OSMI-4 vs. DMSO” = comparison of treatment bead backgrounds (**Fig. S3F**). “LM fOGT_FLAG_” = linear model (LM) background-corrected fOGT_FLAG_ OSMI-4 vs. DMSO AP-MS analysis (Fig. 4B). Purple = increase with OSMI-4; Green = decrease with OSMI-4. Asterisk indicates nominal p-value *p* < 0.05. ns = not significant (nominal *p* > 0.05). **E. CARM1 abundance and OGT interaction status across analyses. “**Original fOGT_FLAG_” = initial fOGT_FLAG_ OSMI-4 vs. DMSO AP-MS analysis (Fig. 3B). “Protein abundance” = average proteome-level abundance (Fig. 3D). “stOGT_FLAG_ OSMI-4 vs. DMSO” = comparison of treatment bead backgrounds (**Fig. S3F**). “LM fOGT_FLAG_” = linear model (LM) background-corrected fOGT_FLAG_ OSMI-4 vs. DMSO AP-MS analysis (Fig. 4B). Purple = increase with OSMI-4; Green = decrease with OSMI-4. Asterisk indicates nominal p-value *p* < 0.05. ns = not significant (nominal *p* > 0.05). **F. Western blot analysis of HCF-1-V5 anti-V5 reciprocal immunoprecipitation (Co-IP).** The Co-IP was performed from HCF-1-V5, 3x-FLAG-OGT mESC whole cell lysate; the control was untagged HCF-1, 3x-FLAG-OGT mESC whole cell lysate. Ratios of OGT levels to HCF-1 levels (OSMI-4, relative to DMSO) in the IP eluate (25% bead boil) and input-normalized ratios were calculated via densitometry analysis in ImageJ. The asterisk indicates bands from the antibody heavy chain.

Overall, this statistical approach eliminated 71% of background binders appearing to increase in interaction and 84% of background binders appearing to decrease in interaction **(Figure 4C)**. This approach also reduced false negatives: the linear model data uncovered 16 proteins significantly increasing in interaction **(Supplementary Figure S4A)** and 38 proteins significantly decreasing in interaction **(Supplementary Figure S4B)**. For example, NUP62 and CARM1 were not shown to differentially interact with OGT in the OSMI-4 vs. DMSO fOGT_FLAG_ pairwise comparison, nor change in abundance. However, they were both significantly enriched in one treatment bead background versus the other, so upon background correction, NUP62 significantly increased **(Figure 4D)** and CARM1 decreased **(Figure 4E)** in interaction with OGT. As for CARM1, these results were anticipated: gene set enrichment analysis (GSEA) terms related to oxidative stress and ROS were enriched in the OSMI-4-treated proteome **(Supplementary Figure S4C)** and it was previously described that CARM1’s interaction with OGT decreases with cellular oxidative stress (59). This showed that this multi-factor modeling strategy can resolve false negatives and correct for factors contributing to experimental variance beyond abundance fluctuations.

We also wanted to validate the finding that HCF-1 does not significantly gain interaction with OGT upon OSMI-4 treatment, as it has been previously shown that OGT associates more with the HCF-1 precursor relative to the cleaved protein (60), and upon OSMI-4 treatment we saw that HCF-1 cleavage was reduced **(Supplementary Figure S4D)**. To do so, we installed a V5 epitope tag on the C-terminus of HCF-1 in the 3x-FLAG-OGT mESC line and then performed an anti-V5 reciprocal immunoprecipitation with or without 20 µM OSMI-4 treatment for 24 hours. Evaluation of HCF-1-associated OGT levels by western blot and densitometry analysis showed that after correcting for increased OGT and decreased HCF-1 levels, HCF-1 does not gain interaction with OGT **(Figure 4F)**. This shows that even after correcting for changes in the background proteome via statistical modeling, non-significant changes in OGT PPIs are maintained and validated through alternative assays.

Beyond known OGT interactors changing in interaction, we also identified proteins which gain interaction with OGT upon its catalytic inhibition that have not previously been described as interactors. While the PR-DUB complex components BAP1 and ASXL1 are well-known OGT interactors (49), the subunit FOXK1 has never been identified in an OGT co-IP. We found that FOXK1 interacts with OGT upon OGT catalytic inhibition, which could indicate that FOXK1 is an indirect or weak interactor of OGT, and with OGT’s increased interaction with BAP1/ASXL1, enough FOXK1 is present to be enriched over baseline. Overall, this exemplifies that by fitting a linear model to enrichment and control samples across conditions, a direct and rigorous comparison between enrichments can be made, allowing for identification of OGT PPIs that change upon OGT catalytic inhibition.

## Discussion

Here, we demonstrate the necessity of correcting for changes in the background proteome in AP-MS experiments that investigate PPI dynamics across different biological contexts. By integrating orthogonal tag controls, parallel proteome-level sample profiling, and multi-factor statistical modeling with traditional AP-MS controls, we confidently elucidate context-dependent PPIs and provide preliminary mechanistic insight, guiding downstream validation and improving data reproducibility.

The use of mock purifications to control for contaminating background binders has been successfully applied for several decades, allowing researchers to avoid harsh purification schemes and incorporate quantitative mass spectrometry techniques (61,62). This study extends this concept to the comparison of affinity enrichments performed from different background proteomes. We anticipate that background proteome correction will become even more crucial as DIA is adopted for AP-MS experiments: greater sampling depths in AP-MS experiments will perturb interactor-level FDR control, affecting one’s ability to properly characterize false positive and negative interactions (8,63). Increased sensitivity will also lead to detection of more condition-specific background binders, which risk misclassification as true interactors by PPI scoring algorithms that rely heavily on bait exclusivity (9,64). To mitigate this, we strongly recommend including controls in which the target protein is present to the same magnitude between the enrichment and control samples but cannot be enriched by the anti-epitope affinity matrix. There may be cases where the endogenous protein of interest cannot be epitope tagged–for example, in post-mitotic or transfection-recalcitrant cells. For this, we suggest engineering two transgene overexpression constructs with distinct epitope tags, rather than relying on a fluorescent protein or empty vector control, to ensure that the control captures any changes to the bead background induced by overexpression of the target protein. Utilizing these tailored controls, as well as having a control per each biologically distinct condition in study, is sufficient and preferred for context-variable PPI characterization. This is because these controls account for all context-dependent factors—not just protein abundance—that may affect a protein’s propensity to be a background binder (*e.g.*, changes in subcellular location, biophysical properties) (65). However, this approach may be infeasible when performing hundreds of AP-MS experiments against different targets in tandem. In these cases, parallel proteome profiling can provide insight as to which proteins may be differential background binders due to abundance changes and thus warrant further validation. Sampling the enrichment input within any experimental design is beneficial because not only does this help explain background proteome differences, but preliminary mechanistic insight is also gained, highlighting whether a protein’s change in interaction may be a direct result of the biological perturbation or through an indirect mechanism leading to its change in abundance and thus interaction stoichiometry.

While this study highlights the importance of background proteome correction for dynamic PPI identification, this concept is also common for other enrichment- and mass spectrometry-based methods. For example, when measuring changes in post-translational modifications (PTMs) between biological contexts, one can also measure global protein abundance via mass spectrometry-based proteomics to determine whether the apparent change is reflective of altered site stoichiometry or of a change in protein abundance (66). Similarly, techniques such as proximity labeling benefit from the use of no-biotin or isogenic controls, which can account for transgene-induced cellular changes (67,68). We therefore expect that background proteome correction will be a necessary consideration for any protein-level, mass spectrometry-based method that utilizes an affinity enrichment matrix.

A limitation of this study, shared with many AP-MS studies (44,69), is that cryptic sources of technical variance affect statistical power. This uncertainty limits one’s ability to truly distinguish true interactors from noise. Several strategies can help address this, including setting stricter significance thresholds, adding more replicates (70), or streamlining sample preparation using standardized protocols and automation (71). However, each of these approaches come with tradeoffs: rote adherence to significance thresholds risks missing true biological changes, while the latter suggestions can be infeasible due to cost or lack of experience, particularly for research groups that do not routinely perform AP-MS experiments. Our experimental framework improves statistical rigor by accounting for variance induced by changes in the background proteome, but further improvements to increase statistical power can be made. In any case, we recommend careful consideration of the model system and target protein biology when prioritizing hits for follow-up studies, and validating differential interactors through orthogonal methods, such as reciprocal immunoprecipitation or proximity ligation assay.

Ultimately, integrating proteome-level information and simple statistical analyses into AP-MS experiments, especially those analyzing PPI dynamics across changing biological contexts, will produce higher quality data in which the rich information and clarity outweigh the additional experimental effort. With advances in the analytical aspects of PPI identification, our experimental paradigms should proportionally progress to ensure data fidelity and reproducibility.

## Data Availability & Supplemental Information

### Data Availability

The original mass spectra files have been uploaded to MassIVE. The Shiny app presented in this work is publicly available on Github (https://github.com/melinabrunelli/LinearModelR).

### Supplemental Data

This article contains supplemental data. Please see “Supplementary Data” for descriptions of supplemental tables and figures.

## Acknowledgements

We thank members of the Myers lab, particularly Sophia Chau, Felipe Vasquez-Castro, and Dominic McGrosso, for their helpful discussions and critical review of the manuscript.

## Funding and Additional Information

This work was supported by NIH NIGMS R35GM147554 and the Global Autoimmune Institute (S.A.M.). S.A.M. is also supported by Anthony R. Carr, Richard S. and Karna S. Bodman, Fred and Pam Wasserman, the Rosemary Kraemer Raitt Foundation Trust, Northern Trust, Christopher and Rebecca Twomey, and Bridget Cresto and the Cresto Family Giving Fund. M.A.B. was supported in part by the UCSD Graduate Training Program in Cellular and Molecular Pharmacology (NIH NIGMS T32 GM007752).

## Author Contributions

M.A.B. and S.A.M. conceptualization; M.A.B., N.M.C., and D.R.M. data curation; M.A.B. formal analysis; S.A.M. funding acquisition; M.A.B. and L.M.C. investigation; M.A.B. and S.A.M. methodology; M.A.B. and S.A.M. project administration; S.A.M. resources; N.M.C. and D.R.M. software; S.A.M. supervision; M.A.B. and S.A.M. validation; M.A.B. visualization; M.A.B. and S.A.M. writing–original draft; M.A.B., L.M.C., N.M.C., D.R.M., and S.A.M. writing–review and editing.

## Abbreviations

AP-MS: affinity purification-mass spectrometry
PPIs: protein-protein interactions
O-GlcNAc: β-N-acetylglucosamine
OGT: O-GlcNAc transferase
mESC: mouse embryonic stem cells
HDR: homology directed repair
nDIA: narrow-window data-independent acquisition

## Supplementary Data

**Supplementary Figure 1.**
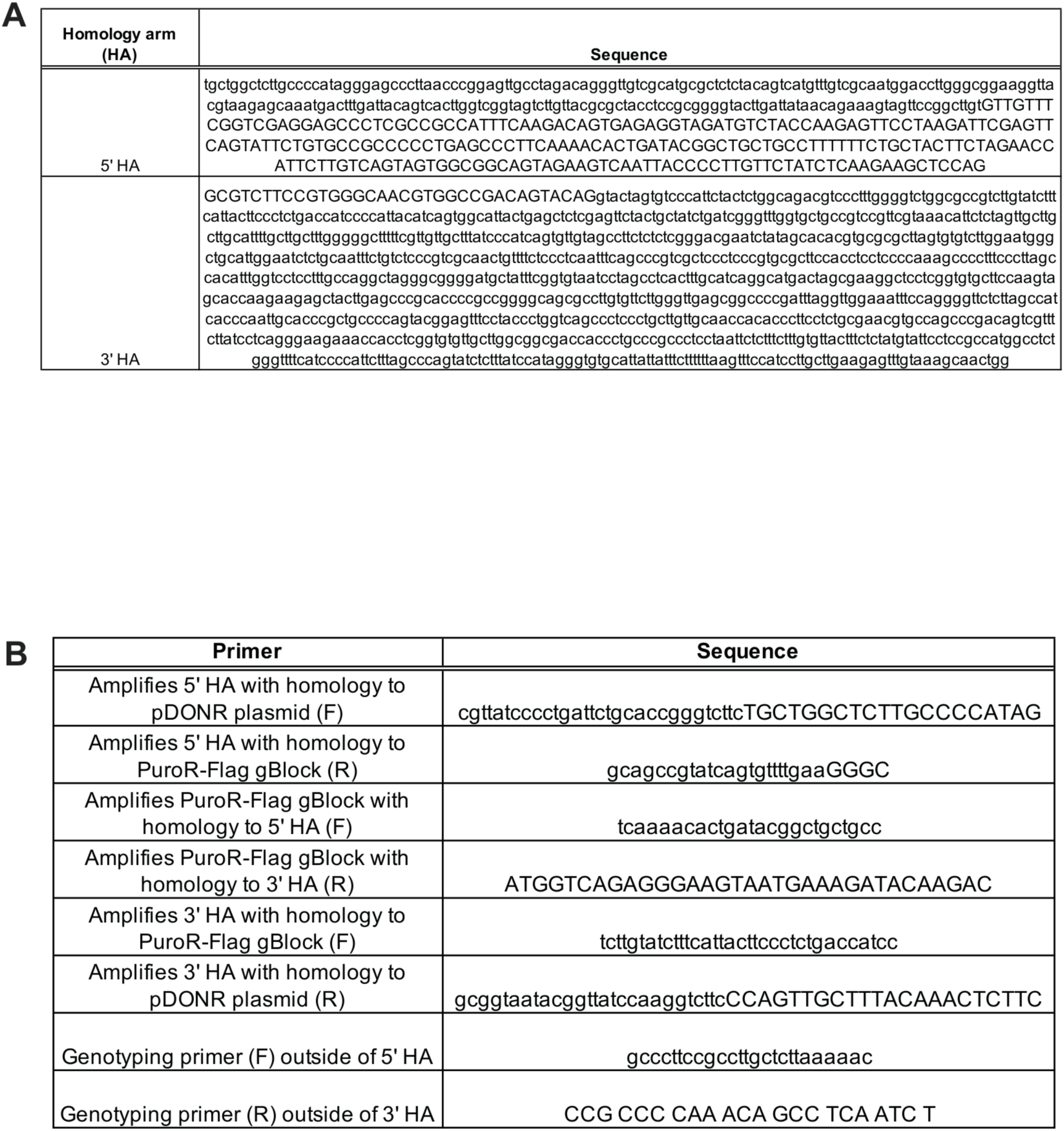
**A.** Table containing sequences of the 5’ and 3’ homology arms (HAs) added to the repair template for knock-in into the endogenous *Ogt* locus. **B.** Table containing primer sequences used to clone the *Ogt* repair template (HAs, PuroR-T2A-3xFLAG or 2x-Strep cassette) into a pDONR-BFP plasmid.

**Supplementary Figure 2.**
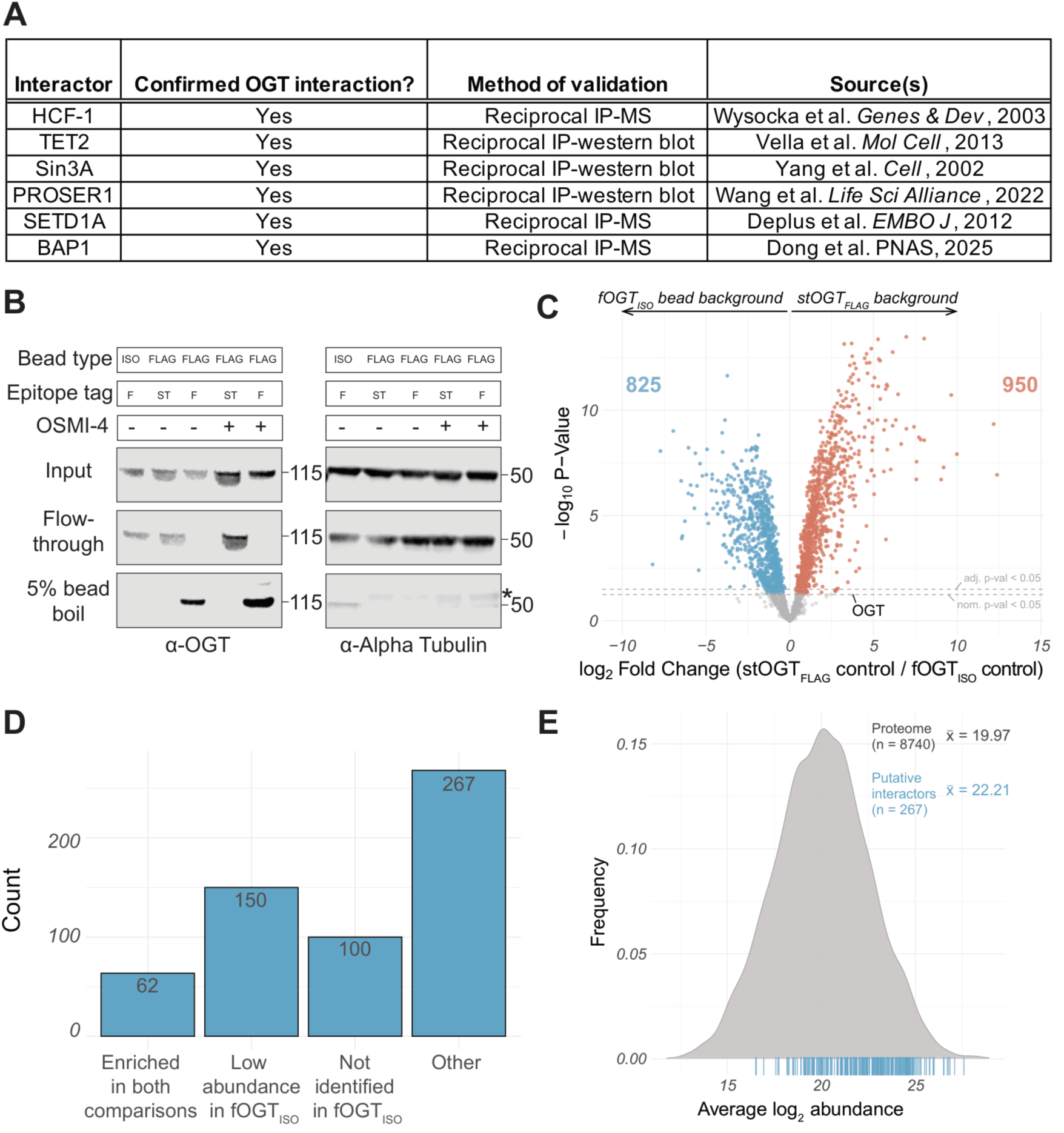
**A. Table containing validated OGT interactors.** A protein is a validated OGT interactor if it has been identified to interact with OGT through some orthogonal method (to OGT co-IP-MS), such as reciprocal immunoprecipitation. **B. Western blot depicting anti-FLAG OGT enrichment efficiency.** Input, flow-through/unbound, and supernatant from 5% boiled beads (eluate) were probed with anti-OGT and anti-Alpha Tubulin antibodies to show FLAG-specific depletion of OGT from mESC lysate and presence on anti-FLAG beads. Asterisk indicates antibody heavy chain. **C. Volcano plot comparing the fOGT_ISO_ and stOGT_FLAG_ controls, an ancillary comparison as an example.** Orange points are proteins specifically enriched in stOGT_FLAG_ samples, and light blue points are proteins specifically enriched in fOGT_ISO_ samples. Gray points have nominal *p* > 0.05. Bolded values specify proteins passing the nominal p-value threshold *p* < 0.05. **D. Bar graph categorizing the 579 putative interactors identified in the fOGT_FLAG_ vs. fOGT_ISO_ comparison.** “Enriched in both comparisons”: log_2_FC > 0 & *p* < 0.05 in the fOGT_FLAG_ vs. fOGT_FLAG_ comparison and fOGT_FLAG_ vs. stOGT_FLAG_ comparisons. “Low abundance”: bottom 20% of abundance in fOGT_ISO_ samples. “Not identified”: proteins not detected in fOGT_ISO_ samples. “Other” encapsulates all leftover putative interactors. **E. Density plot demonstrating abundance bias of “other” putative interactors.** Log_2_ average abundance of proteins in the 3x-FLAG-OGT mESC proteome are plotted. The 274 “Other” proteins from **Fig. S2D** are plotted via a light blue rug plot below the density plot. The x-bar values in the top right corner detail the mean abundance of proteins across the proteome and the mean abundance of the “other” putative interactors.

**Supplementary Figure 3.**
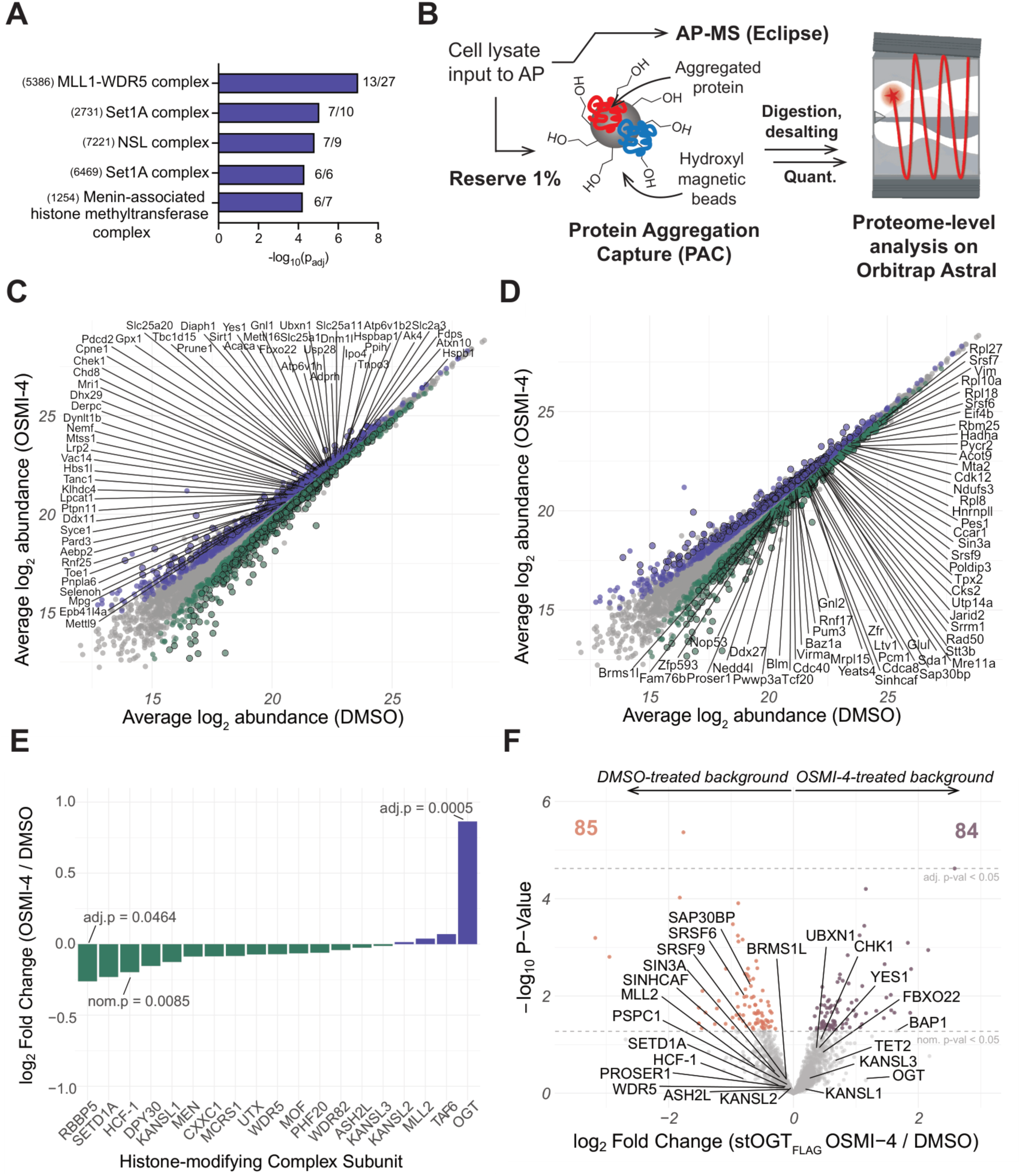
**A. Bar plot of CORUM complex terms relating to the NSL and COMPASS family complexes.** All proteins that significantly increase in interaction with OGT were analyzed in g:Profiler with the following parameters changed from default: species = *Mus musculus*, FDR < 0.01, and no electronic GO annotations. Labels to the right of the bars indicate the number of proteins in the analyzed data set / number of proteins in the annotated CORUM complex. CORUM complex IDs are to the left of the complex name. **B. Schematic of proteome-level sample analysis via LC-MS/MS.** A portion (1%) of the sample input to the affinity purification was reserved as “proteome-level” samples. Protein aggregation capture (PAC) was used to remove contaminants (salts, detergent, cell debris) and allow for on-bead digestion. After MS sample preparation steps, samples were analyzed on an Orbitrap Astral using narrow window data-independent acquisition (nDIA). **C. Scatter plot of proteins** (**56**) **that increase in interaction and abundance upon OSMI-4 treatment.** The average log_2_ normalized protein abundance is plotted (OSMI-4 on y-axis; DMSO on x-axis). Larger dots with a black border are proteins that pass adjusted p-value < 0.05. Purple = interaction gain with OSMI-4; Green = interaction loss with OSMI-4; Gray = nominal *p* > 0.05. Proteins are labeled by their gene symbol. **D. Scatter plot of proteins** (**55**) **that decrease in interaction and abundance upon OSMI-4 treatment.** The average normalized protein abundance is plotted (OSMI-4 on y-axis; DMSO on x-axis). Larger dots with a black border are proteins that pass the threshold adj. p-value < 0.05. Purple = interaction gain with OSMI-4; Green = interaction loss with OSMI-4; Gray = nominal *p* > 0.05. Proteins are labeled by their gene symbol. **E. Bar plot of log_2_ fold changes (OSMI-4 / DMSO) of NSL/COMPASS complex components.** Values are from the OSMI-4-vs. DMSO-treatment proteome-level comparison. Proteins that pass a nominal or adjusted p-value threshold are labeled with their p-value. **F. Volcano plot comparing OSMI-4- and DMSO-treated stOGT_FLAG_ controls, an ancillary comparison as an example.** Proteins enriched in the OSMI-4-treatment background are in mauve; proteins enriched in the DMSO-treatment background are in orange. Bolded values specify proteins passing the nominal p-value threshold *p* < 0.05.

**Supplemental Figure 4.**
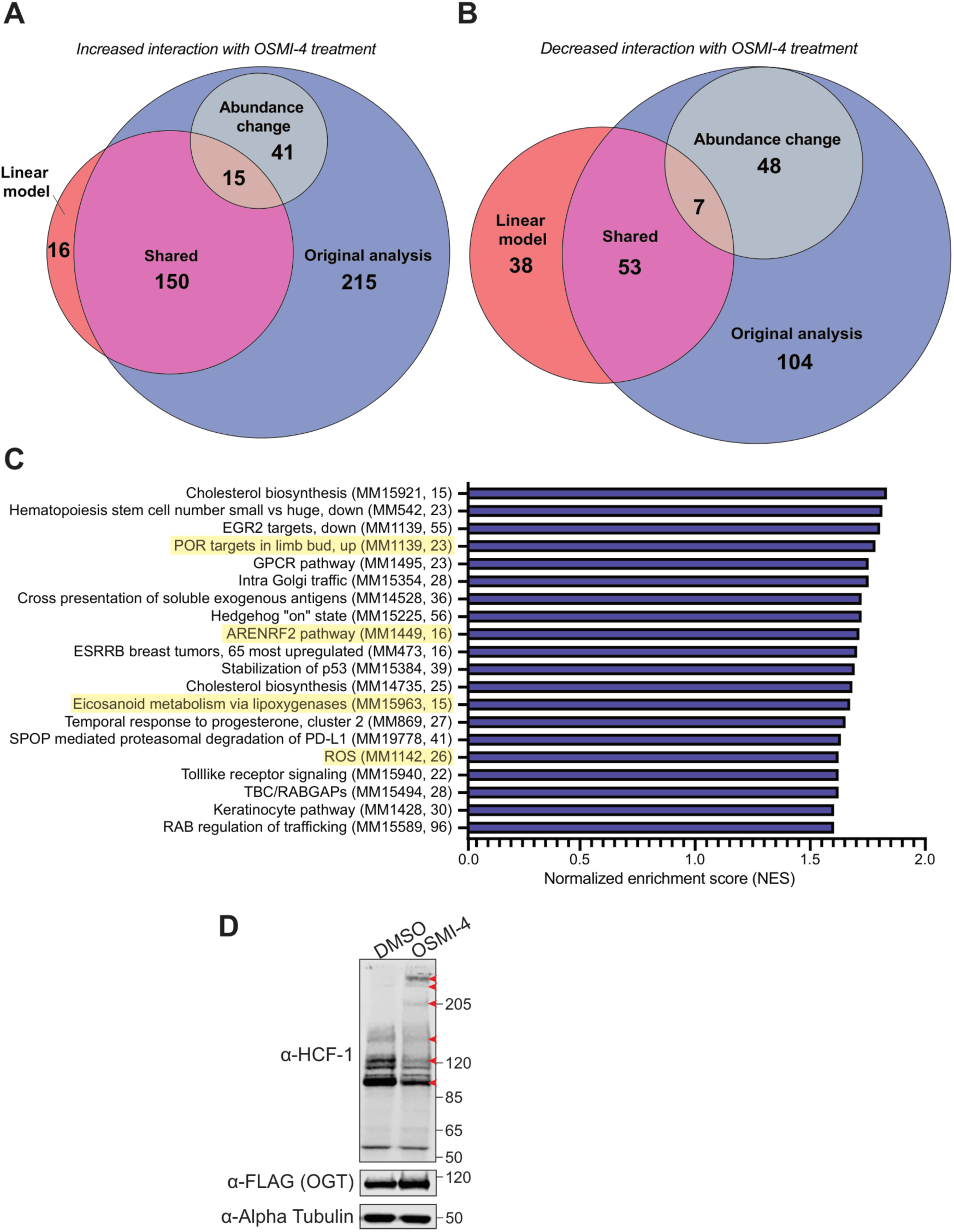
**A. Venn diagram comparing proteins identified by the original OSMI-4 vs. DMSO fOGT_FLAG_ analysis (blue) and the model-fitted analysis (red) that increase in interaction with OGT upon OSMI-4 treatment.** Significant interactors that are identified by both analyses are in pink. Proteins whose abundance increases commensurate with their interaction are in light blue. **B. Venn diagram comparing proteins identified by the original OSMI-4 vs. DMSO fOGT_FLAG_ analysis (blue) and the model-fitted analysis (red) that decrease in interaction with OGT upon OSMI-4 treatment.** Significant interactors that are identified by both analyses are in pink. Proteins whose abundance decreases commensurate with their interaction are in light blue. **C. Normalized enrichment scores (NES) of a GSEA comparing the OSMI-4 and DMSO-treated proteomes.** The top 20 enriched gene set terms are shown. Yellow highlighted terms are related to oxidative stress or reactive oxygen species (ROS) production. The enrichment value used for GSEA analysis was log_2_ fold change. All gene sets pass a nominal p-value threshold *p* < 0.05. The parentheses next to the gene set name specify the mSigDB identifier and the number of genes (proteins) present in the gene set. **D. Western blot analysis of HCF-1 cleavage patterns upon OSMI-4 treatment.** Lysate was run on a 3-8% Tris-Acetate protein gel for SDS-PAGE prior to western blot. The HCF-1 antibody was raised specifically against the HCF-1 C-terminal half. Red arrowheads indicate cleavage products that either increase or decrease in abundance upon OSMI-4 treatment.

### Supplementary Data Tables

**Supplementary Table 1.** nDIA proteomics analysis of 3x-FLAG-OGT and 2x-Strep-OGT mESC lines

**Supplementary Table 2.** AP-MS analysis comparing fOGT_FLAG_, fOGT_ISO_, and stOGT_FLAG_ enrichments

**Supplementary Table 3.** AP-MS analysis of OSMI-4-vs. DMSO-treated fOGT_FLAG_

**Supplementary Table 4.** nDIA proteomics analysis of 3x-FLAG-OGT mESC OSMI-4-vs. DMSO-treated proteomes

**Supplementary Table 5.** Linear model-fitted AP-MS analysis of OSMI-4-vs. DMSO-treated fOGT_FLAG_ and stOGT_FLAG_

## Notes

### Competing Interest Statement

The authors have declared no competing interest.

